# Representation and inference of size control laws by neural network aided point processes

**DOI:** 10.1101/2021.01.24.428011

**Authors:** Atsushi Kamimura, Tetsuya J. Kobayashi

## Abstract

The regulation and coordination of cell growth and division is a long-standing problem in cell physiology. Recent single-cell measurements using microfluidic devices provide quantitative time-series data of various physiological parameters of cells. To clarify the regulatory laws and associated relevant parameters such as cell size, mathematical models have been constructed based on physical insights over the phenomena and tested by their capabilities to reproduce the measured data. However, such a conventional model construction by abduction faces a constant risk that we may overlook important parameters and factors especially when complicated time series data is concerned. In addition, comparing a model and data for validation is not trivial when we work on noisy multi-dimensional data. Using cell size control as an example, we demonstrate that this problem can be addressed by employing a neural network (NN) method, originally developed for history-dependent temporal point processes. The NN can effectively segregate history-dependent deterministic factors and unexplainable noise from a given data by flexibly representing functional forms of the deterministic relation and noise distribution. With this method, we represent and infer birth and division cell size distributions of bacteria and fission yeast. The known size control mechanisms such as adder model are revealed as the conditional dependence of the size distributions on history and their Markovian properties are shown sufficient. In addition, the inferred NN model provides a better data representation for the abductive model searching than descriptive statistics. Thus, the NN method can work as a powerful tool to process the noisy data for uncovering hidden dynamic laws.

## I. INTRODUCTION

One of the major quests in microbial physiology is to unveil the fundamental principles and laws underlying the regulation and coordination of cell growth and division[1]. Recent developments in microfluidic devices enable us to track microbial cells over hundreds of generations[2–5]. Various physiological parameters, in particular cell sizes, have been measured over time quantitatively at the single-cell level [2, 6–8].

To elucidate laws of cell growth and division, a large body of research has been conducted by employing techniques from physical and mathematical sciences[1, 9–14]. Especially, simple physical and mathematical models have been pursued, which can explain cell growth and division with a small number of relevant variables. In order to account for the significant cell-to-cell variety of division patterns in measured data, the process of cell growth and division was typically formalized as a continuous-time stochastic process where each cell grows and divides with a variable rate[10, 11]. In this formalism, different models can be implemented by choosing the relevant variables to determination of the division rate. For example, ‘sizer’ model[10] posits that the absolute size of the cell is relevant whereas ‘adder’ model[15–17] considers the added size from the birth of the cell fundamental. Even though the biological mechanisms regulating cell divisions are complex and can involve many biochemical reactions and signaling, such simple models were found to reproduce the division patterns successfully. However, construction of a good model generally requires us for deep insights and intuitions into the cell physiology as well as trial and error because one has to figure out relevant variables and their relation for characterizing and reproducing cell divisions statistics. Even if an obtained model works good, we may find a better model by choosing or including another variable that we did not try. Moreover, biological data shows high variability, only part of which a simple model can explain. Thus, the unexplained part is represented as noise by assuming its distribution. However, the prefixed repertoire of noise distributions such as Gaussian or exponential distribution may not appropriately explain the unexplained components. Thus, the conventional model construction by abduction faces a constant risk that we may overlook important parameters, variables, and relations especially when the complicated high dimensional or noisy time-series data is concerned and when the model that we are fitting to data does not have a sufficient representation power over both deterministic relation and noise distribution.

In the last decade, machine learning (ML) methods have been prodigiously developed and applied to a wide variety of problems[18]. ML methods especially deep learning (DL) have demonstrated that they can semi-automatically extract complicated patterns underlying large amounts of noisy data sets by removing irrelevant dimensions and unexplainable noise factors. Even though ML, in reality, is not at all the magic wand that can solve problems without any human help or preconceived assumptions, we can use it to support us for searching a defined but huge model space without relying on our insights and intuition.

In this work, we employ a neural network (NN) method to the problem how cell size is regulated. By following the similar formulation in the previous modeling of cell size control as stochastic process[10, 11], we represent the cell size dynamics interrupted by division events by using temporal point process (TPP). Then we introduce a NN method to represent the intensity function of the process, which can depend flexibly on the history of cell size over the retrospective lineage. By training the NN, we obtain conditional probability density functions (PDFs) of cell sizes at birth and division, given their past histories. The trained PDFs reproduce and confirm the previous results and presumed assumptions of size control; the cell division is indeed stochastic; the Markovian model is sufficient for predicting birth and division sizes. Adder and weak-sizer principles for *E. coli* and *S. pombe* respectively are obtained from the history dependency of the conditional PDFs. Moreover, the NN method is shown to extract the relations between multiple variables and provide a better data representation for the model searching by abduction than the conventional descriptive data plotting or use of summary statistics. Thus, the NN method can work as a powerful tool for uncovering hidden dynamic laws from noisy data.

The rest of the paper is organized as follows. In Section II, we extend the formalism of the stochastic process of cell growth and division to the case where cell sizes at division depend on their history. We then explain the formulation and recent developments of intensity-based NN models for TPP. In Section III, we describe the experimental data of *E. coli* and *S. pombe* cell division used in our analysis. In Section IV, we first present performances of existing intensity-based models and show that a fully NN model developed recently is the best in its flexible expressiveness of data. By using the model, we examine how the conditional PDFs of cell sizes depend on their history. We also examine the ‘typical’ behavior by calculating the medians of the conditional PDFs and compare them with the size control laws obtained by previous modeling. In Section V, we summarize and discuss our study.

## II. FORMULATION

### A. Modeling history-dependent stochastic dynamics of cell size

The stochastic dynamics of cell size can be modeled as a history-dependent temporal point process (hTTP) [19] First, we characterize the history of a cell by a sequence of cell sizes at birth (*l*_*b*_) and division (*l*_*d*_). *m* is the pre-determined length of the sequence with alternating *l*_*b*_ and *l*_*d*_. We categorize sequences into two types ***h***_*m*_ and ***g***_*m*_: sequence ***h***_*m*_ is the size history up to the most recent birth *l*_*b*_ and ***g***_*m*_ is that up to the most recent division *l*_*d*_. For even *m*, they are represented as

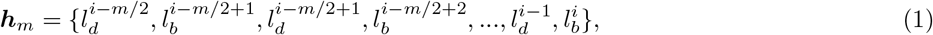

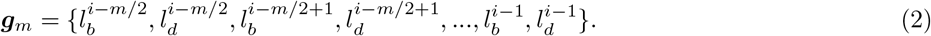

and for odd *m*,

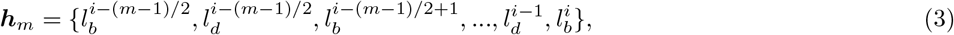

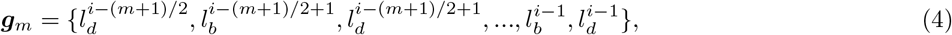

where the superscripts, e.g., *i*, denote the generations of the cell defined along a lineage.

Then, we consider the next division size 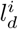 of a cell with history ***h***_*m*_. The history includes sizes up to the most recent birth size 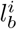. We define the survival probability Π(*l*|***h***_*m*_) that the cell with the history ***h***_*m*_ does not divide up to size *l*. In the following, we assume that the size dynamics is stationary and thus Π(*l*|***h***_*m*_) is independent of generation *i*. The survival probability satisfies the following master equation:

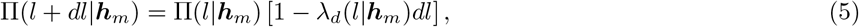

where *λ*_*d*_(*l*|***h***_*m*_)*dl* is the probability that division occurs within the size interval [*l, l* + *dl*] given the size history ***h***_*m*_ and the fact that the cell did not divide up to size *l*. In the continuum limit, Eq. (5) is written as

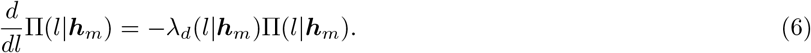

The formal solution of Eq. (6) is

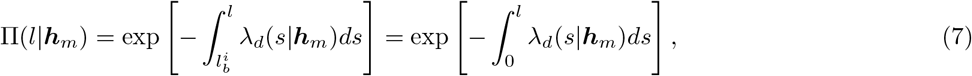

at the second equality of which, the integral range is changed by defining *λ*_*d*_(*s*|***h***_*m*_) = 0 for 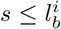. Since the probability that a cell division occurs within the size interval [*l, l* + *dl*] is

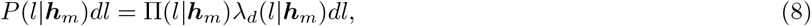

the probability density function (PDF), *P* (*l*|***h***_*m*_), of division-size conditioned by history is written as

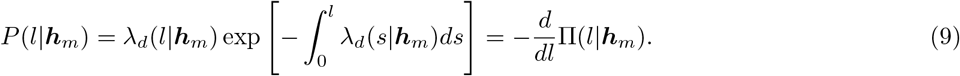

In a similar manner, one can describe the birth-size PDF, *Q*(*l*|***g***_*m*_), for 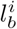:

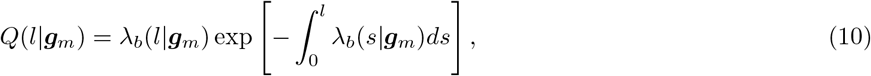

where the history ***g***_*m*_ includes the division size 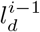 up to the generation *i* − 1. The stationarity for generation is also assumed. Even though the rate function *λ*_*b*_(*l*|***g***_*m*_)*dl* itself does not have an appropriate physical interpretation for birth size, we can use *λ*_*b*_(*l*|***g***_*m*_) in modeling as a proxy of *Q*(*l*|***g***_*m*_) because *λ*_*b*_(*l*|***g***_*m*_) has the same information as *Q*(*l*|***g***_*m*_).

### B. Representation and inference of temporal point process by neural networks

Temporal point process (TTP) is a stochastic process that describes a sequence of discrete events at times 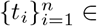 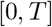. The history-dependent conditional intensity function *λ*(*t*|***h***_*t*_) *≥* 0 is typically used to specify the dependency of the next event time *t* on the event timing history ***h***_*t*_ = {*t*_*i*_ : *t*_*i*_ < *t*}.

Given the conditional intensity function, one can obtain the conditional PDF of the next event time *t*_*i+1*_ as

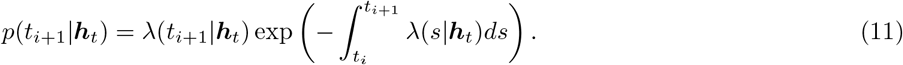

While *λ*(*t*_*i+1*_|***h***_*t*_) is an indirectly way to represent *p*(*t*_*i+1*_|***h***_*t*_), we need not care about the constraint ∫ *p*(*t*_*i+1*_|***h***_*t*_)d*t*_*i+1*_ = 1 if we use *λ*(*t*_*i+1*_|***h***_*t*_). In the conventional point process modeling and inference, simple functional forms have been assumed for *λ*(*t*_*i+1*_|***h***_*t*_; ***θ***) or *p*(*t*_*i+1*_|***h***_*t*_; ***θ***) to make their parameter inference of ***θ*** and log-likehood computation tractable. However, the expression power of the model is severely restricted and insufficient especially when *λ*(*t*|***h***) should be a complicated function of either *t* or history ***h*** or both.

In the last couple of years, neural networks have been employed in different ways. For example, a recurrent neural network (RNN) was used to obtain a fixed-dimensional representation ***r***_*t*_, which compresses the information of history ***h***_*t*_[20]. Then, ***r***_*t*_ is used with a simple and fixed form of *λ*(*t*|***r***_*t*_; ***θ***).

The representation power was extended further by employing another NN to flexibly model the functional form of *λ*(*t*|***r***_*t*_; ***θ***). Among others, Omi et.al.[21] proposed a fully NN model, in which the integral 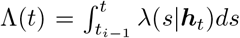 rather than *λ*(*s*|***h***_*t*_) is modeled by a feed-forward NN. This allows us to efficiently compute the log-likelihood, 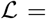 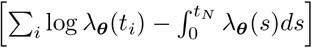, by avoiding integration of *λ*(*s*|***h***_*t*_), regardless of the functional form of the intensity function *λ*(*s*|***h***_*t*_). This method endows more flexiblity in the hTPP modeling, reduces computational complexity for parameter learning, and thereby extends its applicability.

These NN models can be directly applied to the size control of cell physiology because the mathematical framework is almost identical: the conditional PDF in Eq. (11) has the same form with Eq. (9) and Eq. (10). In this paper, we apply FullyNN model to obtain the conditional PDFs of division and birth sizes on size history.

## III. DATA

In this study, we use two public data sets[8, 22]. Both were obtained by microfluidic single-cell measurement devices, versions of the commonly known ‘mother machine’[2, 6]. The devices allow tracking of mother cells trapped at the bottom of the observation channels over tens or a hundred of generations (see Fig. 1A). From the data obtained, one can extract the following parameters to characterize the size dynamics and divisions events (see Fig. 1B): birth size *l*_*b*_, division size *l*_*d*_, the size added between a consecutive pair of birth and division Δ_*d*_ = *l*_*d*_−*l*_*b*_, the relative septum position *l*_1/2_, and division interval *τ*.

**FIG. 1.**
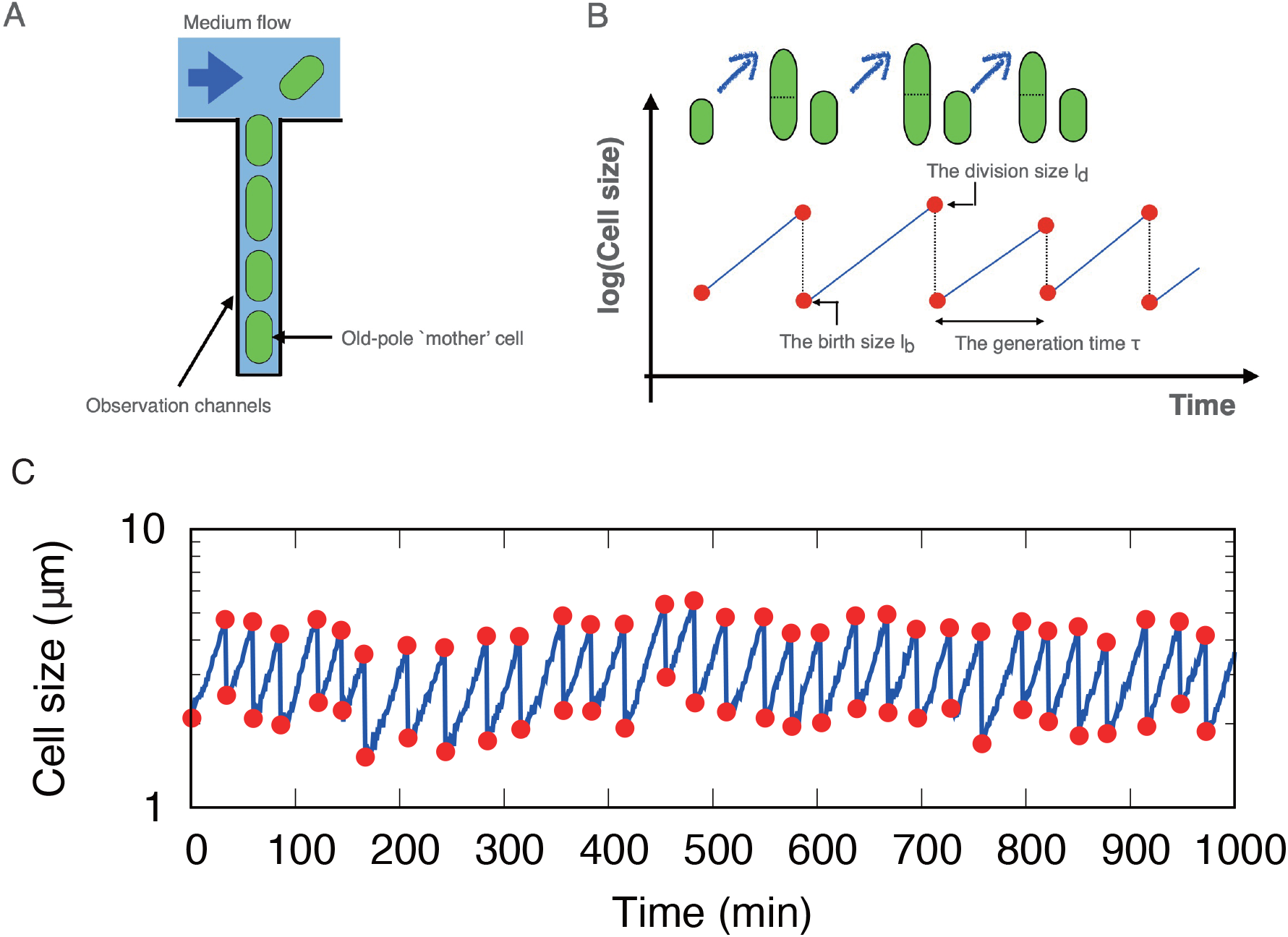
(A) Schematic illustration of the microfluidic device. The old-pole mother cell is trapped at the bottom of the observation channel. (B) Definition of the size parameters. Δ_*d*_ = *l*_*d*_ − *l*_*b*_ is defined by birth and division sizes of the same generation (cell cycle). The relative septum position *l*_1*/*2_ is defined by 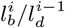, where *i* − 1 and *i* denote indices of consecutive generations. (C) An example of cell size trajectory from the data of *E. coli* at 37°C.

The first data set was obtained by the measurements of *Escherichia coli* (*E. coli*)[22]. The data were recorded every minute and contain time series of cell length of mother cells. The measurements were conducted in three different growth conditions of temperatures (37°C, 27°C, and 25°C). An example of the cell size time course is shown in Fig. 1C.

The second set was obtained by the measurements of *Schizosaccharomyces pombe* (*S. pombe*)[8]. The data were recorded every three minutes and contain time series of cell area. The measurements were conducted in seven different culture conditions with different media and temperatures (28°C, 30°C, and 34°C in yeast extract medium [YE], and 28°C, 30°C, 32°C and 34°C in Edinburgh minimum medium [EMM]).

In Appendices A and B, we describe the details of preparing and preprocessing the sequential data and procedures for NN learning.

Basic statistics of the parameters are summarized in Tables S1, S2 and S3 of the supplementary materials for *E.coli* and *S. pombe* in YE and EMM conditions, respectively. We also present the histograms of cell sizes in Fig. S1, S2 and S3 of the supplementary materials for *E.coli* and *S. pombe* in YE and EMM conditions, respectively.

## IV. RESULTS

### A. The fully NN model achieves a better performance than the other RNN based models

In this section, we first test performances of different RNN methods by applying them to the cell-size data of *E. coli*. and *S. pombe*. In the previous study [21], FullyNN model we use in our subsequent analysis achieved a competitive or better performance over other RNN based models for various synthetic and real data. Here, we will show that this is also the case for the cell-size data.

In addition to FullyNN model, we here consider three RNN models with specific intensity functions of *λ*_*b*_ and *λ*_*d*_. The simplest one is to assume a constant intensity function (constant model) *λ*\*(*l*_*i*_) = exp(***v***^*T*^***h***_*i*_ + *b*), where ***h***_*i*_ denotes the embedded vector of the history by RNN, and ***v***^*T*^ and *b* are learnable parameters. This constant intensity function corresponds to an exponential distribution for the PDF in Eq. (11), whose parameter varies over generations depending on the history ***h***_*i*_. The second one is the exponential intensity function (exponential model) *λ*\*(*l*_*i*_) = exp(*wl*_*i*_ + ***v***^T^ ***h***_i_ + *b*) where it depends on the size *l* but can be integrated directly[20]. This exponential intensity function corresponds to a Gompertz distribution for the conditional PDF. As a more flexible functional form[23], a piecewise constant model is considered where the intensity function is discretized by piecewise constant functions as 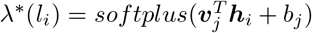 for (*j*−1)*L* ≤ *l* ≤ *jL*, where *j* = 1, 2, …, *l*_*max*_/*L* with given *l*_*max*_ and *L*. In this study, we fix *L* = 128 and 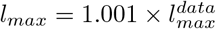, where 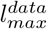 denotes the maximum size in the data.

Figures 2A and B show examples of trajectories of the actual observations and the RNN models after training for birth and division sizes of *E. coli*, respectively. For each RNN model and each generation *i*, the median of the PDF is calculated by using the observed history of sizes before *i*. The medians of the FullyNN and the piecewise constant models agree with the actual observations better than the other models. Significant deviations from the measured data were observed in the exponential model for birth size and in the constant model for both sizes.

**FIG. 2.**
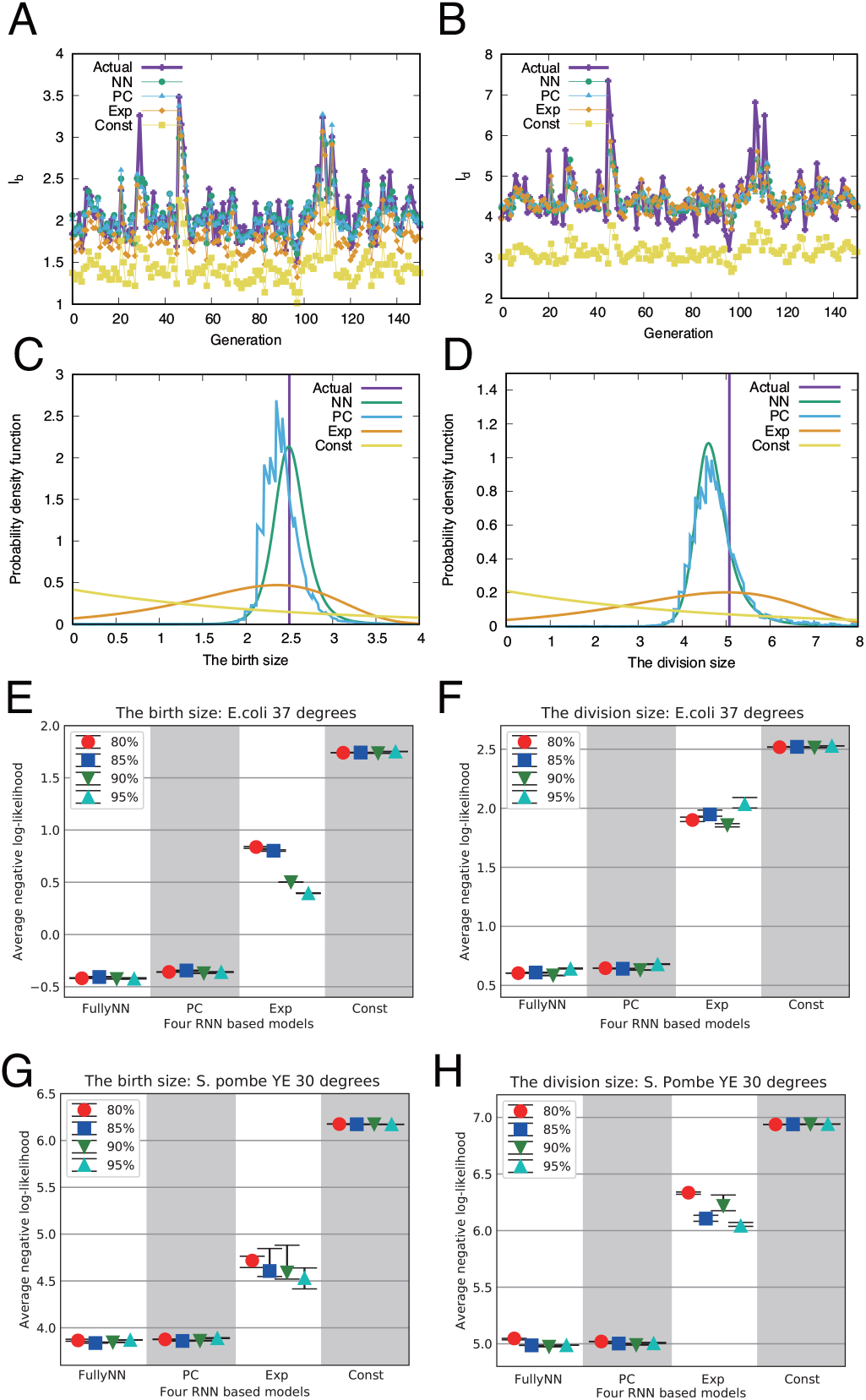
(A,B) Trajectories of the actual observations and the medians of conditional probability density functions (PDFs) of for *E. coli* at 37°C. The PDFs are estimated by the four RNN based models for (A) birth size and (B) division size. (C,D) The shapes of the PDFs of (C) birth size and (D) division size at the generation *i* = 30. (E, F, G, H) The performances of the four models for (E) birth size and (F) division size of *E. coli* at 37°C, and for (G) birth size and (H) division size of *S. pombe* in YE condition at 30°C. The performances of the four RNN based models are evaluated by the negative log-likelihood. The log-likelihood is averaged over the test data. Lower score means a better predictive power. The history length *m* is set to 10.

In Figs. 2CD, we calculate the shapes of the PDFs for the four models at generation *i* = 30 in Figs. 2AB. It should be noted that the PDFs are history-dependent and thereby their shapes change depending on the generation at which we calculate them. The conditional PDFs of the FullyNN and the piecewise constant models are located relatively close to the actual observation and their shapes are similar. The PDFs of the exponential and the constant models are much broader and their functional forms are restricted to the Gompertz and exponential functions, respectively.

To measure the performances of the four models quantitatively, we calculated negative log-likehood 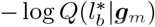 and 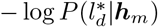 for birth and division sizes, respectively, where 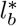 and 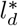 denote the actual observations in the validation data. A lower negative log-likehood means a better performance in prediction. Figures 2E-H show average performances of the four models for both *E. coli* and *S. pombe*. For almost all cases, the FullyNN model achieves the best performance with the smallest values of the negative log-likelihood. Since the amount of data is smaller than those used in Ref. [21], we change the fraction of the training data from 80% to 95%. We found that the scores are approximately constant for the different sizes of training data, which assures that the FullyNN model is sufficiently trained. The piecewise constant model performs similarly well but slightly worse than the FullyNN model and predicts rugged conditional PDFs.

The exponential and constant models perform poorly because they have the restrictions in the functional form. The PDF of cell sizes is typically single peaked and right-skewed (see section IV C). Therefore, they cannot be approximated sufficiently by a Gompertz or an exponential distribution. Also, the performances of the exponential model are unstable. Its scores are more sensitive to the size of training data than the others. In any cases, the FullyNN model performs better by flexibly approximating the PDFs.

### B. The most recent size in history is sufficient for birth and division size prediction

The NN method has several hyperparameters. Of particular interest is the length *m* of the histories ***h***_*m*_ and ***g***_*m*_. In the previous modeling of size control, Markov models were employed and their sufficiency were not tested. Only recently, the impact of the length of history was tested by using a regression model[24]. The dependence of the performance on the history length *m* can be used to systematically and quantitatively verify how far back in the history is relevant to predict the next division or birth size.

Figure 3 shows the performance of the FullyNN model for *l*_*b*_ and *l*_*d*_ as functions of history length *m*. Even though a slight improvement between *m* = 1 and *m* = 2 is observed for the division size *l*_*d*_ of *E. coli.* (Fig. 3B), the performances do not show significant improvements with increasing *m* for *m* ≥ 1. This observation indicates that the most recent elements in the history (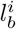 in ***h***_*m*_ and 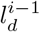 in ***g***_*m*_ are practically sufficient. This supports the assumptions of the previous size control models.

**FIG. 3.**
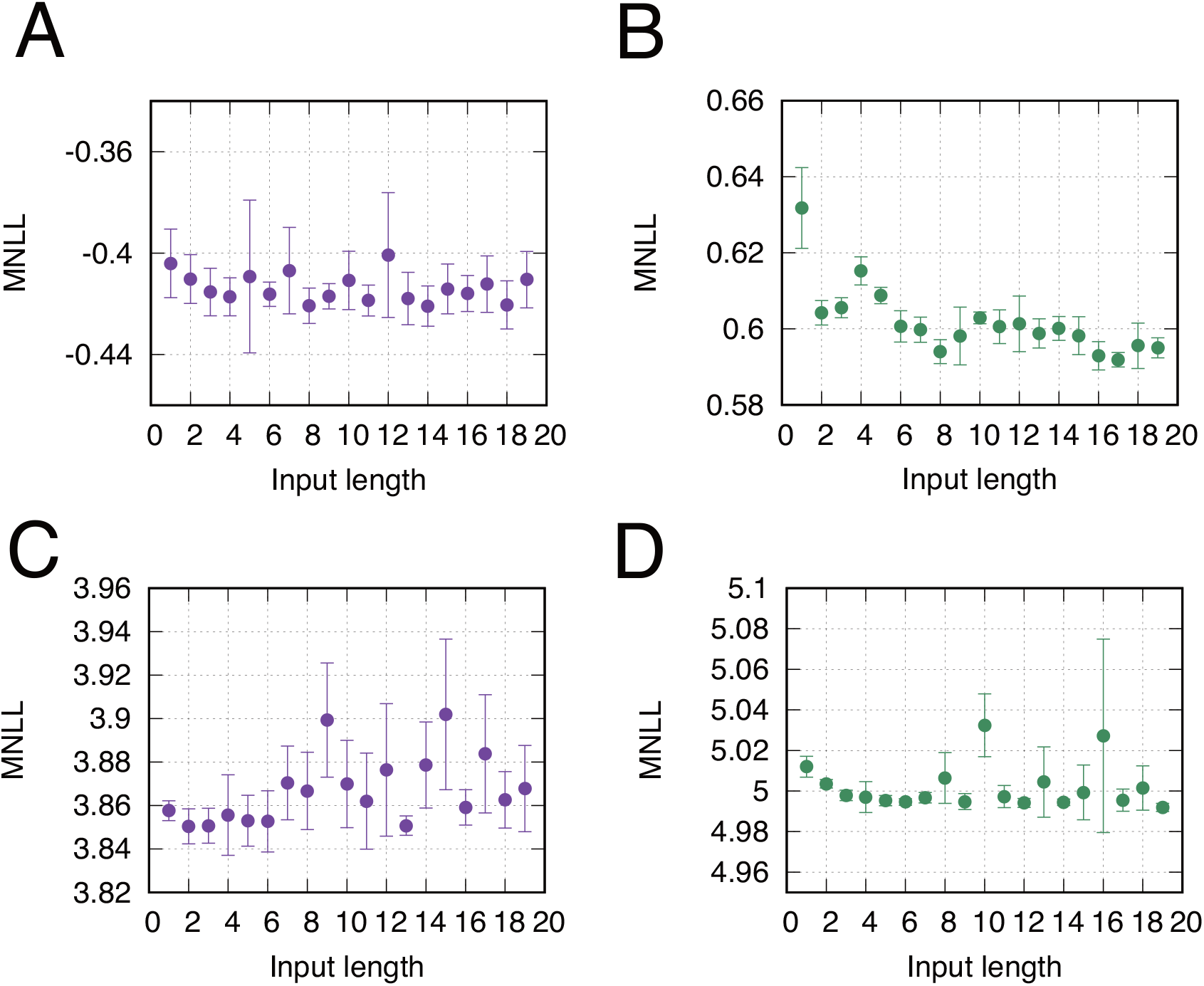
Performances with different history length *m* for (A) birth size and (B) division size of *E. coli* at 37°C, and for (C) birth size and (D) division size of *S. pombe* in YE condition at 30°C. See Fig. S4 of the supplementary materials for *S. pombe* in EMM condition. The performances are evaluated by the negative log-likelihood. The log-likelihood is averaged over the test data.

We also investigated the performance as functions of other three NN hyperparameters, the number of layers and the number of units in each layer (Fig. S5 and S6 in the supplementary materials) to confirm that increase in the complexity of NN from current parameter values does not significantly improve the results.

### C. Shapes of birth and division distributions depend differently on history

We next analyze how the shapes of the birth and division size distributions depends on history by using the estimated PDFs, *P* (*l*_*d*_|***h***_*m*_) and *Q*(*l*_*b*_|***g***_*m*_).

Figure 4AB shows the PDF of *E. coli* birth size *Q*(*l*|***g***_*m*_). While the shape of the PDF can in principle depends on all the elements in the history ***g***_*m*_, we exclusively focus on the most recent division size because we verified it as the most influential and sufficient element of the history in Fig. 3 (see also Fig. S7 and S8 in the supplementary materials for the dependence of *P* (*l*|***h***_*m*_) and *Q*(*l*|***g***_*m*_) on the earlier sizes).

**FIG. 4.**
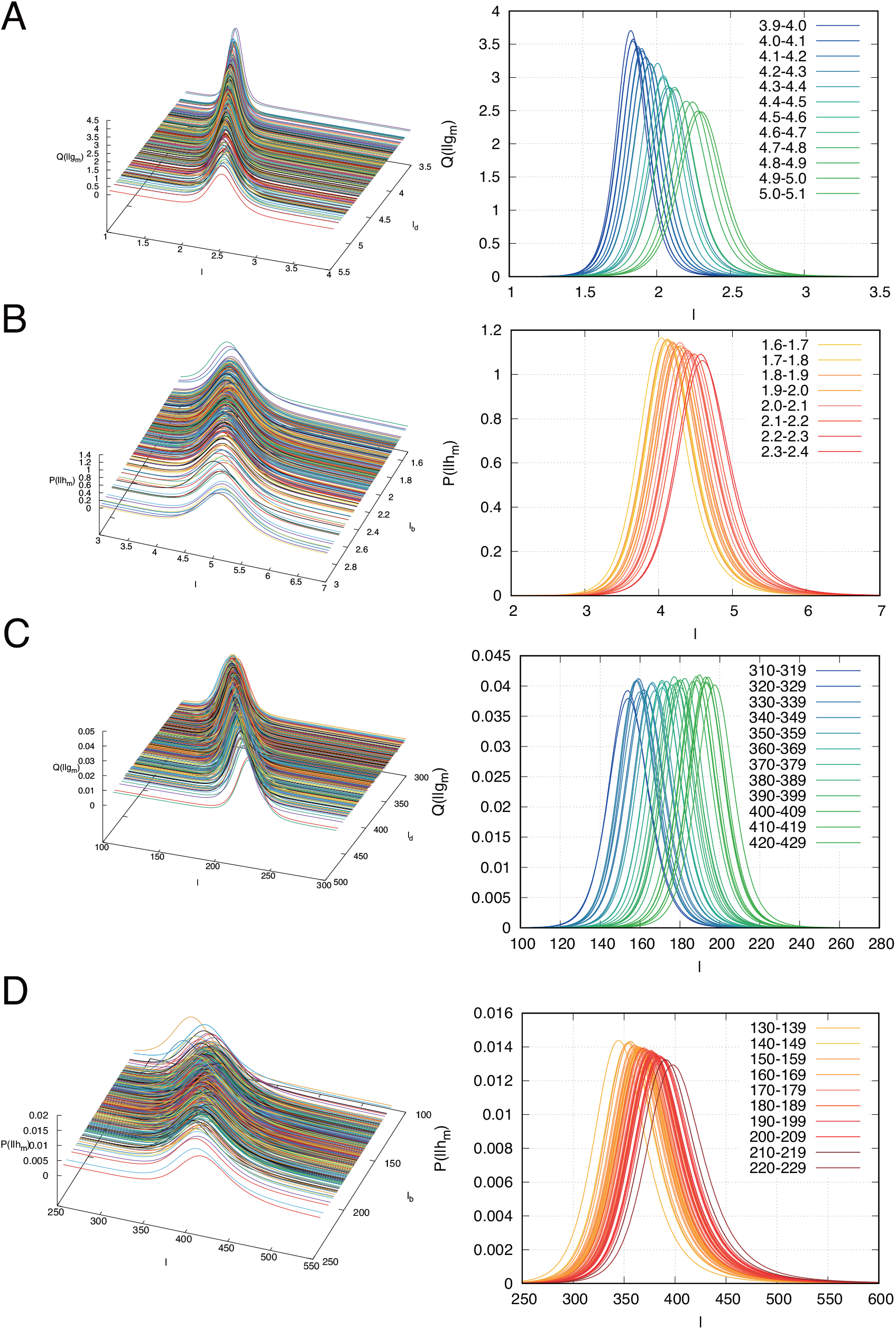
The conditional probability density functions (PDFs) for (A) birth size *Q*(*l*|***g***_*m*_) and (B) division size *P* (*l*|***h***_*m*_) of *E. coli* at 37°C, and those for (C) birth size and (D) division size of *S. pombe* in YE condition at 30°C. *Q*(*l*|***g***_*m*_) and *P* (*l*|***h***_*m*_) are plotted as a function of the most recent size, i.e., *l*_*d*_ for *Q*(*l*|***g***_*m*_) and *l*_*b*_ for *P* (*l*|***h***_*m*_). Left panels are 3D plots of the PDFs as functions of *l* and the most recent size in the histories. Right panels are the slices of the PDFs for fixed values of the most recent size. In right panels, the colors of the curves indicate a range of the most recent size. One can see that the sliced PDFs shift from left to right as the most recent size increases. The history length is set to *m* = 10.

As demonstrated in Fig. 4A, the mode of the birth size PDF is monotonously dependent on the most recent division size, which reflects the obvious fact that a bigger cell divides into two bigger daughters on average. The modes of *Q*(*l*|***g***_*m*_) is approximately located at the position *l* = *l*_*d*_ × *l*_1/2_, where the relative septum position 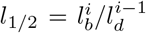 is approximately 0.46 for this data set (see Table S1 in the supplementary materials). This dependency of birth size PDF is captured even when only the most recent division size is considered as the history, i.e., by setting *m* = 1 in the model when trained (see Fig. S9 in the supplementary materials).

Similar dependency of mode is observed for the division size PDF *P*(*l*|***h***_*m*_) as a function of the most recent birth size (Fig. 4B). This dependency, however, is less prominent than that of the birth size PDF, suggesting that the division size is under tighter control than the birth size.

In addition, Fig. 4A clarifies that the birth size of a cell with a bigger parent cell shows greater variation than one with a smaller parent. This is a sharp contrast to the birth size PDF of *S. pombe*. (Fig. 4C for YE and Fig. S10 for EMM condition), whose variation, i.e., the width of the PDF, is almost independent of the previous division size. This results may indicate the difference in the birth size control between *E. coli* and *S. pombe.*, the latter of which has a more elaborated molecular mechanism to determine the middle of a cell [25, 26]. For the mode of division size PDF of *S. pombe*. in Fig. 4D, in contrast, we observe a similar dependency on the birth size to *E. coli*, which implies the existence of division size control for both *E. coli* and *S. pombe*.

All these properties are automatically extracted and clearly presented in the estimated conditional PDFs by the Fully NN model from the noisy raw data.

### D. NN makes adder principle conspicuous by denoising data

We further elucidate the “typical” behaviors of birth and division size distributions as functions of histories, by calculating the medians of the PDFs estimated by the FullyNN model. We have a stable and systematic way to obtain the median of the PDF represented implicitly by the complicated NNs (see Material and Method). Figure 5A exemplifies a comparison of the time-series of actual birth *l*_*b*_ and division *l*_*d*_ sizes with the medians of the PDFs for *E. coli* (the same data with Fig. 2AB).

**FIG. 5.**
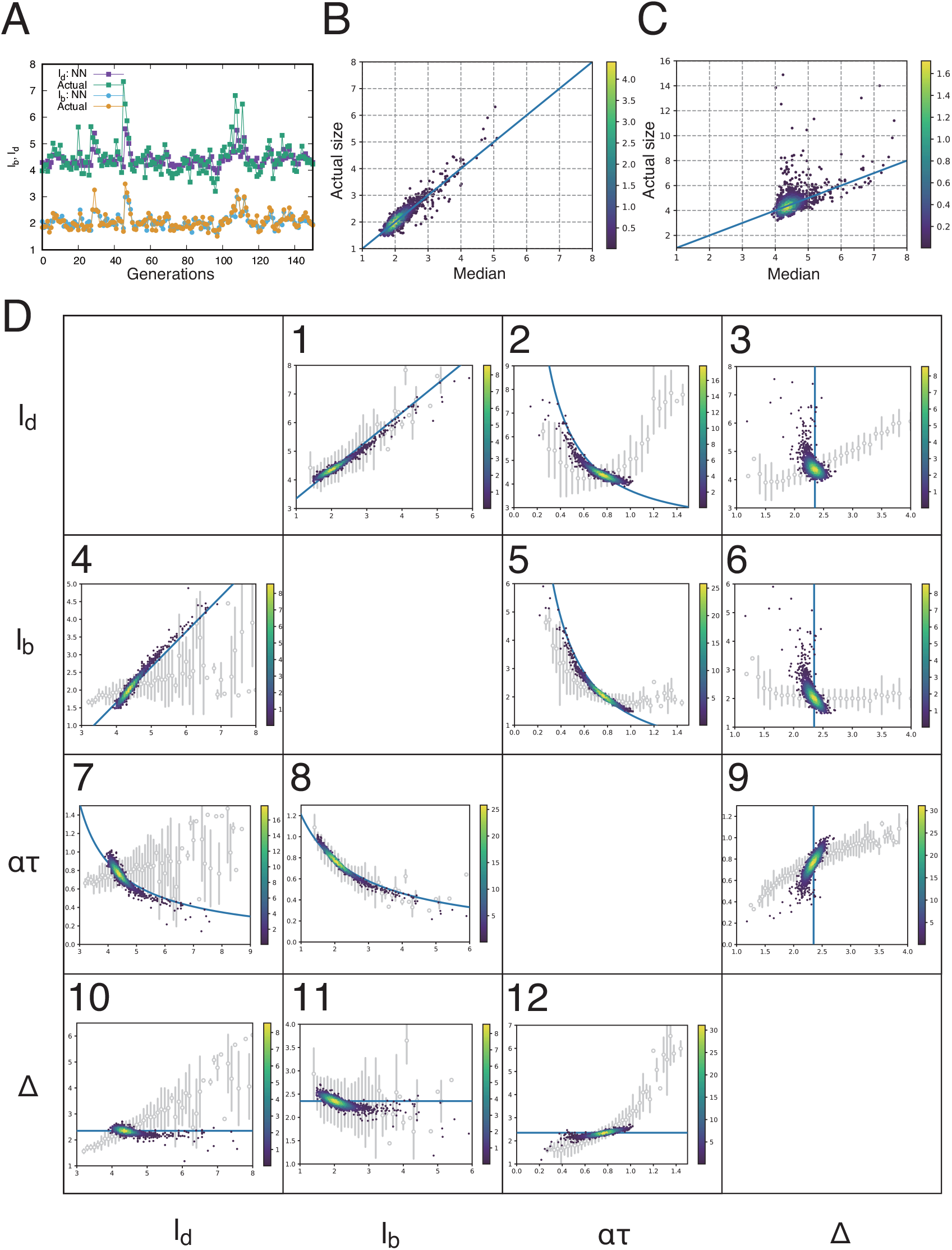
Data for *E. coli.* at 37°C. (A) Trajectories of the actual birth and division sizes and the medians of the PDFs obtained by the NN model. (B, C) Density plots of the actual sizes and the medians of the PDFs for (B) birth size and for (C) division size. The identity lines are also shown in blue. (D) All correlations between all pairs of the four parameters, *l*_*d*_, *l*_*b*_, *ατ*, Δ. Each density plot shows the correlation between each pair of the parameters calculated from the medians of PDFs. Given the birth size of *l*_*b*_, the PDF for the division size is used to calculate the division size (*l*_*d*_), elongation *ατ* = log(*l*_*d*_/*l*_*b*_), and the added size Δ = *l*_*d*_−*l*_*b*_. The blue curves indicate the adder model with Δ = 2.35. The gray empty circles and error bars indicate the summary statistics of the binned raw data. The circles and bars are the medians in the binned range and their interquartile range, respectively.

For birth size *l*_*b*_, the actual observation (orange) and the medians (light blue) agree well and are distributed along *x* = *y* in the scatter plot (Fig. 5B). This result demonstrates that the median of the PDF works as a good predictor of *l*_*b*_. For the division size *l*_*d*_ in Fig. 5A, we observe more sporadic jumps in the actual observations, which the median of *P* (*l*|***h***_*m*_) (purple) fails to predict. Except such outliers, e.g., *l* > 10, the median can predict a general trend in the division size over time, which is also verified in the scatter plot of Fig.5C.

By using the PDF, we next try to detect the adder law of *E. coli* size control revealed by the previous modeling. It posits that cells add constant size Δ between birth and division, irrespective of the birth size. A standard approach to explore size control laws is to plot correlations among pairs of potential control parameters. Fig. 5D shows the correlations between four parameters, division size *l*_*d*_, birth size *l*_*b*_, elongation *ατ*, and added size Δ for *E. coli* at 37°C.

We first focus on the correlation between the division size *l*_*d*_ and the birth size *l*_*b*_ (Fig. 5D1). The plot clearly show a positive correlation in which a cell with a large (small) birth size divides at a large (small) division size. The trend is also consistent with binned raw data indicated by empty circles and error bars, each of which corresponds to the median and the interquartile range of the binned data, respectively.

The correlation between the added size Δ = *l*_*d*_−*l*_*b*_ and the birth size *l*_*b*_ shows a slightly negative correlation for 37°C (Fig. 5D11) and approximately no correlation for 27°C and 25°C (Fig. S11 and S12 in the supplementary materials). This negative correlation is dimmed in the binned raw data (empty circles). Owing to the reduced variation by the NN model, we can identify the weak sizer property in 5D 11 at 37°C. Overall, these results support the adder principle even though it is not necessarily perfect at high temperature.

### E. Underlying relations may not be represented appropriately by a simple descriptive approach

To investigate whether adder principle can account for all the other correlations, we also present the expected behaviors of the ideal adder model by the blue curves in Fig. 5D (see section 1 in the supplementary materials for the ideal adder). For most of plots in Fig. 5D, the medians agree very well with the idealized adder model (the blue curves).

In contrast, in all the plots except Figs. 5D 1, 8, and 11, the binned raw data fail to capture the correlations expected from the adder model. It should be noted that the binned plots basically represent the actual distributions of data (Fig. S13 of the supplementary material). The deviation are not produced by the binning procedure except the non-monotonical behavior between elongation and division size (Fig.5D2) where the data size is relatively small for small *ατ* s. The discrepancies of the binned data arise because substantial variations are present in the actual data, and the noise simultaneously influences multiple variables so that an additional correlation is produced between the parameters. Thus, when we do not know the relevant variables *a priori*, we can obtain a correct relation only if we happen to choose a right variable as the binning variable as in 5D 1 and 8. Otherwise, the binned data may mislead us over the underlying relation. By using the conditional PDF and its representative value, the effect of additional correlations is eliminated between fluctuating variables and a direct comparison becomes possible between data and models.

However, there is still a possibility that the discrepancy between the median and the binned data is an artifact of the NN modeling. Actually, the median does not perfectly follow the idealized adder relation in some plots such as 5D 3, 6, and 9. To exclude this possibility, we constructed a synthetic data of a stochastic adder model, in which the added size is generated from a fixed right-skewed distribution by reflecting the right-skewness in the observed division distribution (see Table S4 and section 2 in the supplementary material). Then, we trained the NN model with this synthetic data (Fig. 6). Here, we assume that the birth size is given by a perfect binary division 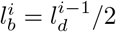 for simplicity, and the division size is given by 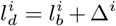, where the added size Δ^*i*^ is drawn from a log-normal distribution.

**FIG. 6.**
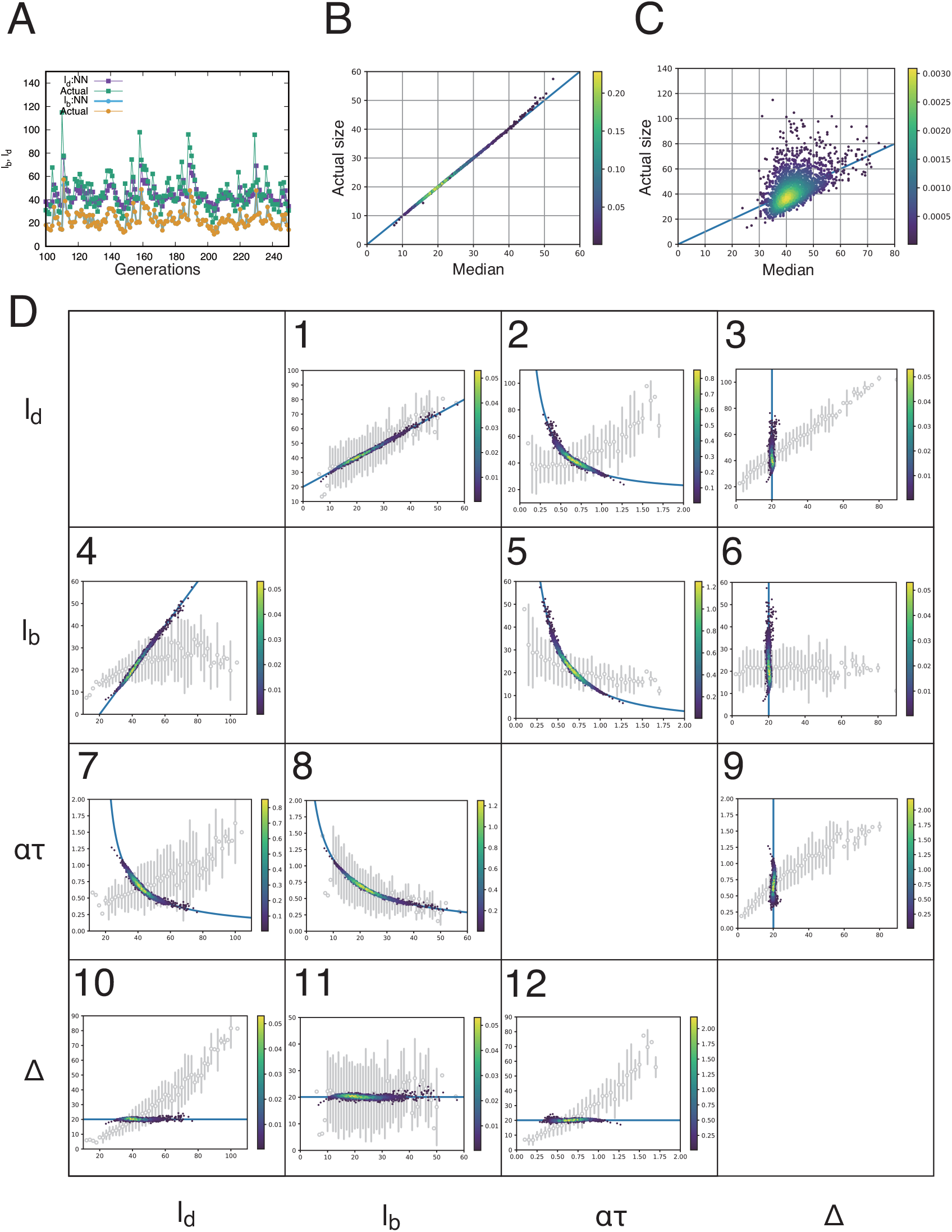
Data for synthetic data of the adder model. The alternating sequence of *l*_*b*_ and *l*_*d*_ is synthesized as follows. For the birth size, a perfect binary division is assumed so that 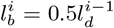 where the superscript denotes the generation. For the division size 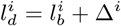 the added size is drawn from the log-normal distribution *L*(*μ, σ*^2^) with *μ* = 3 and *σ* = 0.5. (A) Trajectories of the actual birth and division sizes and the medians of the PDFs obtained by the NN model. (B, C) Density plots of the actual sizes and the medians of the PDFs for (B) birth size and for (C) division size. The identity lines are also shown in blue. (D) All correlations between all pairs of the four parameters, *l*_*d*_, *l*_*b*_, *ατ*, Δ. Each density plot shows the correlation between each pair of the parameters calculated from the medians of PDFs. Given the birth size of *l*_*b*_, the PDF for the division size is used to calculate the division size (*l*_*d*_), elongation *ατ* = log(*l*_*d*_/*l*_*b*_), and the added size Δ = *l*_*d*_−*l*_*b*_. The blue curves indicate the adder model with Δ = exp(*μ*). The gray empty circles and error bars indicate the summary statistics of the binned raw data. The circles and bars are the medians in the binned range and their interquartile range, respectively.

For the synthetic data, the birth size is predicted almost perfectly by median of the trained conditional PDF (Fig. 6B). In addition, Fig. 6C of the synthetic data reproduced an asymmetry between the median and the actual size in Fig. 5C such that the points distribute around the line *y* = *x* but larger deviations appear more above the line. Thus, this asymmetry comes from the right-skewness of the added size and thereby of division size, which is not sufficiently represented only by the median of the PDF. Figure 6 D also confirms that the median of PDF can perfectly reproduce the assumed adder property whereas the binned plots (gray empty circles and error bars) fail to capture the property except in Fig. 6 D 1, 8, and 11, similarly to the case of *E. coli* data.

These results highlight a general advantage of the NN model over the conventional descriptive plot of raw data especially when we do not know which variables should work as explanatory variables in advance.

In this sense, the NN model can also benefit the physical modeling by abduction by extracting relations among multiple variables from noisy data.

### F. NN works for other microbe with different size control mechanism

Finally, we investigate *S. pombe*, whose size control law differs from the perfect adder [17, 27].

Figure 7 shows the same analysis as that in Fig. 5 for YE condition at 30°C (see also Fig. S14 for EMM condition). The elongation is omitted because the growth of *S. pombe* may depart from exponential trajectory and the equation *l*_*d*_ = *l*_*b*_ exp(*ατ*) used for *E. coli* may be invalid (see Fig. S15).

**FIG. 7.**
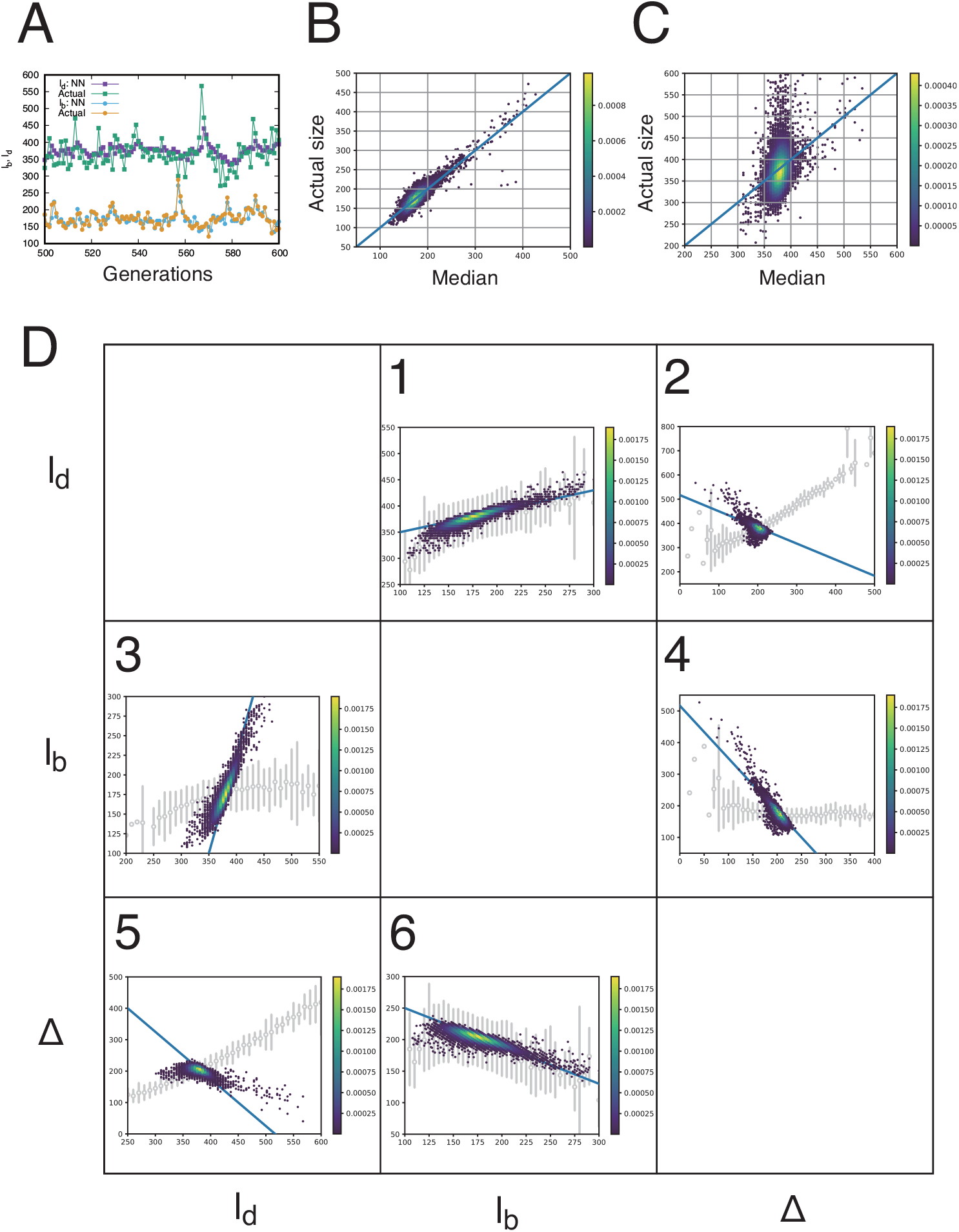
Data for *S. pombe* in YE condition at 30°C. (A) Trajectories of the actual birth and division sizes and the medians of the PDFs obtained by the NN model. (B, C) Density plots of the actual sizes and the medians of the PDFs for (B) birth size and for (C) division size. The identity lines are also shown in blue. (D) All correlations between all pairs of the three parameters, *l*_*d*_, *l*_*b*_, Δ. Each density plot shows the correlation between each pair of the parameters calculated from the medians of PDFs. Given the birth size of *l*_*b*_, the PDF for the division size is used to calculate the division size (*l*_*d*_) and the added size Δ = *l*_*d*_−*l*_*b*_. The blue lines indicate Δ = −0.6*l*_*b*_ + 310. The gray empty circles and error bars indicate the summary statistics of the binned raw data. The circles and bars are the medians in the binned range and their interquartile range, respectively.

The NN model reproduces the birth size of *S. pombe* as good as that of *E. coli* (Fig. 7 A and B). In addition, Fig. 7D6 clarifies a significant negative correlation between the birth size and the added size, whose slope is approximately −0.6 (the blue line). The slope of −1 in this plot posits the perfect sizer principle in which there is a critical size for division. The approximated slope of −0.6 indicates a size control in-between perfect adder and sizer, which is consistent with previous studies [17, 27].

Similarly to Fig. 5D, all the plots of median in Fig. 7D agree with the weak-sizer model of −0.6 (blue curves). The binned data fails to capture the relations except in Figs. 5D 1 and 6. Thus, this result reinforces the general advantage of the NN method illustrated with *E. coli* data.

By comparing the results of *E. coli* and *S. pombe*, one noticeable difference is observed between Figs. 5C and 7C, where the actual sizes of *S. pombe* deviate from the prediction by medians. This deviation can be attributed to the sizer aspect of *S. pombe* size control. The perfect sizer principle posits that the division occurs at a certain threshold size, which makes the average division size independent of the birth size as well as the other elements in the size history. Thus, if size control becomes closer to sizer, the best statistical model is the history-independent iid model that faithfully reproduce the variation of division size around the threshold size.

To order to confirm this, we make a weak-sizer model where the added size weakly depends on the birth size and is generated in a probabilistic manner (Fig. 8 and section 2 in the supplementary material). Here, we assume that the birth size is given by a perfect binary division 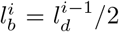, and the division size is given by 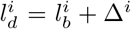 where the added size Δ^*i*^ is a realization drawn from a normal distribution with the mean (median) −0.6*l*_*b*_ + 800. Here, we chose the slope −0.6 from Fig. 7D6. With this synthetic data, as shown in Fig. 8C, the NN model reproduces the division size distribution similar to that in Fig. 7C. Also, the other plots in Fig. 8 perfectly reproduced the assumed weak-sizer property. Therefore, the NN model can effectively extract the relation between variables hidden by noise and its correlation among different variables.

**FIG. 8.**
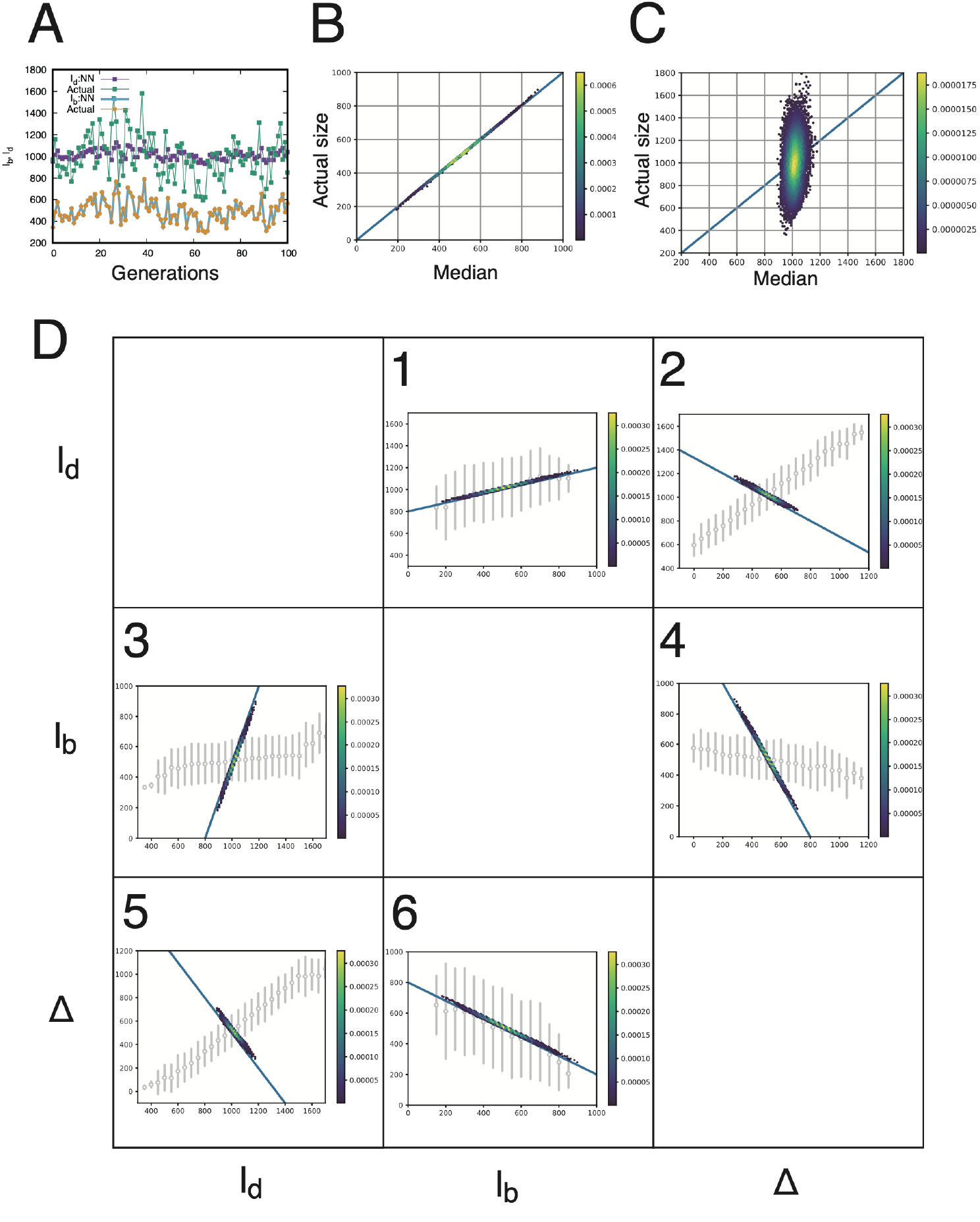
Data for synthetic data of the weak sizer model. The alternating sequence of *l*_*b*_ and *l*_*d*_ is synthesized as follows. For the birth size, a perfect binary division is assumed so that 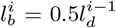, where the superscript denotes the generation. For the division size 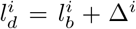, the added size is drawn from the normal distribution *N* (*μ, σ*^2^) with 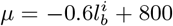 and *σ* = 200. (A) Trajectories of the actual birth and division sizes and the medians of the PDFs obtained by the NN model. (B, C) Density plots of the actual sizes and the medians of the PDFs for (B) birth size and for (C) division size. The identity lines are also shown in blue. (D) All correlations between all pairs of the three parameters, *l*_*d*_, *l*_*b*_, Δ. Each density plot shows the correlation between each pair of the parameters calculated from the medians of PDFs. Given the birth size of *l*_*b*_, the PDF for the division size is used to calculate the division size (*l*_*d*_) and the added size Δ = *l*_*d*_|*l*_*b*_. The blue lines indicate the weak sizer model with *μ* = −0.6*l*_*b*_ + 800. The gray empty circles and error bars indicate the summary statistics of the binned raw data. The circles and bars are the medians in the binned range and their interquartile range, respectively.

## V. CONCLUSION AND DISCUSSION

In this work, we have applied the NN method to the problem of cell size control. Our method describes the size dynamics as a history-dependent temporal point process and models the integral of its intensity function directly by NNs. It is free from assuming any specific functional shape for the size distribution or its dependence on the past size history. Thereby, our method enjoys an extremely flexible expressive power, whose performance was confirmed by using the time-series data of *E.coli.* and *S.pombe.*

One notable advantage of this method is that it can automatically separate two factors in size determination: one is the history-dependent deterministic size control; the other is the stochastic component which cannot be explained only by the size history. From the former factor, we confirmed that the size control can be well approximated as Markovian, which supports the general assumption, presumed in previous modeling, that the birth size is essential for the next division size. From the latter factor, we can know that there still remains a large stochastic component that cannot be explained only by the information of past size, no matter how far back the history is considered. The result implies that the division is determined not only by size but with the interference of other processes in self-replication of a cell.

If we have a substantial portion of stochasticity, which cannot be explained by past sizes, it hampers the discovery of unknown relations from data by abduction[12, 28]. The NN method flexibly represents the stochaticity in data in the form of a probability distribution and its typical behavior as its representative value. This representation allows us to find underlying relationships between variables that are difficult to capture clearly in the conventional scatter plotting or binning of data. The size control relations obtained by NN are consistent with the adder model of *E.coli* and capture the sizer aspect in *Spombe*.

These results demonstrate that the NN mehod is an extremely powerful tool to semi-automatically separate deterministic and stochastic factors from measured data, and it can develop further as an alternative to the conventional modeling method by abduction.

The NN model we used was originally developed for history-dependent time point processes[21]. Therefore, the present method is also directly applicable to cell cycle duration, i.e., division interval. The division interval is another phenotypic variable influenced by and interfering with cell-size control. Similarly to size, the division intervals of mother and daughter cells shows correlations. While models of cell-size control typically predict negative correlations[1, 16], the correlation of division intervals can be positive in a subset of bacterial experiments and most observations of mammalian cells[29]. The positive correlation suggests that the division interval is a heritable quantity over generations, whose dynamics was recently inferred as a latent state dynamics from cellular lineage trees[30]. The NN method we employed may contribute to disentangling the relation between cell size and division interval.

In addition, we may extend its architecture to incorporate cell size, division interval, and also other variables such as gene expression as a multidimensional history. The flexible expression power of the NN method is indispensable for elucidating the complicated mechanisms of cell division where various factors are involved. However, we should mention that, when we incorporate multiple factors, the problem of causality have to be addressed. In the formulation of the point process, a division event is the objective variable whereas other variables including past division events are treated as explanatory variables. When only size is concerned as in this work, the causal relationship between the objective and explanatory variables is obvious. When other factors are involved, however, the true causal relationship among them is not known *a prior*, e.g. we should distinguish whether a high gene expression before division induces cell division with a long division interval or the long division interval leads the high gene expression. In order to address such problems, it would be important to develop a NN method that can integrate causal inference, data from intervention experiments, and other techniques such as the dual reporter system.

## ACKNOWLEDGEMENT

We thank Yuichi Wakamoto, Yuki Sughiyama, So Nakashima, and Dimitri Loutchko for fruitful discussions. This work is supported by JST, CREST Grant Number JPMJCR1927 and JPMJCR2011 and by JSPS KAKENHI Grant Number 19H03216 and 19H05799.

## APPENDIX A Preparation of data

The birth size *l*_*b*_, division size *l*_*d*_, and division interval *τ* are defined from the *E. coli.* data[22] as follows. The data were recorded every minute, and the division events were indicated by ‘division flags’ (1 if division occurred). The birth size *l*_*b*_ is defined by the cell length at which each division occurred, and the division size *l*_*d*_ is defined by the length at the measurement immediately before division. The division interval *τ* is defined by the time between the consecutive division flags.

The measurement was conducted for each cell over 70 generations, i.e., 70 *l*_*b*_s, and 69 *l*_*d*_s and *τ*s. The data were combined for all the measured cells in each of the three growth conditions (37°C, 27°C, 25°C) to assure sufficient training samples for learning. It contains a total of 160, 54 and 65 mother cell lineages at 37°C, 27°C and 25°C, respectively.

For *S. pombe*, we used data published in Ref. [8]. The data were recorded every three minutes and contained millions of time slices. Here, the authors introduced and defined several indices in each time slice to handle the huge data as follows. See Ref. [8] for further details. The status of a cell (alive or dead) was indicated by an index ‘LastIndex’. If the index is 0, the cell was alive. If the index is 1, it survived at the end of the tracking in which an index ‘NextCell’ was set to 0. If the LastIndex was set to 2 or greater, the cell was dead or disappeared from the channel. The division events were indicated by indices ‘MergeIndex’ and ‘NewBornCell’. If the index MergeIndex was set to 1, the cell would undergo cell division by the next time slice. If the index NewBornCell was set to 1, the cell divided between the current and the immediately before time slices.

The parameters for *S. pombe* are defined by monitoring these indices as follows. The birth size *l*_*b*_ is defined by the cell area at which LastIndex ≤ 1 and NewBornCell = 1. The division size *l*_*d*_ is defined by the cell area at which LastIndex ≤ 1 and MergeIndex = 1. The division interval *τ* is defined by the time between the consecutive realizations of LastIndex ≤ 1 and MergeIndex = 1. Here, the time is not calculated if the end of tracking occurred (NextCell = 1) between the realizations.

For each of seven culture conditions with different medium (yeast extract medium [YE] and Edinburgh minimum medium [EMM]) and temperatures, a time window was introduced in the paper within which stable growth was achieved (for example, *t* ≥ 3000(min) in YE conditions). To select the data in the time window, we monitor an index ‘Slice’ in addition to the above indices. For example, the parameters were defined only if the slice *i* satisfied *i* ≥ 1000 for YE conditions where the slice *i* corresponds to *t* = 3(*i* − 1) (min).

## APPENDIX B Training and validation of neural network model

Both for the data of *E. coli.* and *S. pombe*, the sequences of the birth size *l*_*b*_ and the division size *l*_*d*_ were divided into training and validation data. Unless otherwise mentioned, the first 80% and the last 20% of the sequences were used for training and validation, respectively. In the training phase, the model parameters were estimated using the training data. For the optimization, the Adam optimizer with a learning rate of 0.001 was used and the batch size was 256[31].

We performed the training of *l*_*b*_ and *l*_*d*_ separately as follows. For the division size *l*_*d*_, the sequences of *l*_*b*_ and *l*_*d*_ are formatted into the form of ***h***_*m*_ in Eq. (1) and (3) for a given length *m*. To calculate the PDF *P* (*l*|***h***_*m*_) of the next division size of *l* = *l*_*d*_, the model parameters was estimated by using the collection of sequences ***h*_*m*_**s. In the same way, the sequences of *l*_*b*_ and *l*_*d*_ were formatted into the form of ***g***_*m*_ in Eq. (2) and (4) to calculate the PDF *Q*(*l*|***g***_*m*_) of the birth size *l*_*b*_.

In the validation phase, we evaluated the performance of the trained models, respectively for *l*_*b*_ and *l*_*d*_, using the validation data. For each generation time *i*, the PDF of the division size *l*_*d*_ was calculated from Eq. (9) and scored by the negative log-likelihood (NLL), 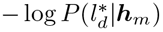, for the actual observation 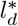 at *i*. Here, a smaller score indicates a better predictive performance for the validation data. Finally, the average of NLLs (MNLL) was calculated for *l*_*d*_s at different generations within the validation data. In the same way, the performance was scored for the birth size of *l*_*b*_. The training and validation of the NN explained above was basically performed using the code provided by Omi et.al. [21], while necessary modifications were made, in particular, to distinguish ***h*_*m*_** and ***g*_*m*_**. The code uses TensorFlow 2.0.0.

Unless otherwise mentioned, the hyperparameters of the NN model are fixed as follows. The length of histories is *m* = 10. In the feed-forward network to learn the integral of the intensity function, the number of units in each layer was 64, and the number of layers was 2. The number of units in RNN to embed the history was fixed to 64.

In order to characterize typical behavior of trained PDFs, we calculated their medians by following Omi et.al.[21]. The median 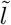 was calculated by solving 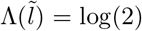, where Λ(*l*) denotes the integral of the intensity function. We also calculate the characteristics in size control models as follows: Given the most recent actual birth size *l*_*b*_ in the history ***h***_***m***_, the added size is calculated as 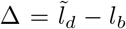 The elongation 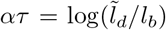 is also computed from the median. Here, we assume the equation *l*_*d*_ = *l*_*b*_ exp(*ατ*) because the majority of *E. coli.* cells elongate exponentially with time and do not display significant growth rate variations at specific cell stages (see Fig. S16 in the supplementary materials).

## Notes

### Competing Interest Statement

The authors have declared no competing interest.

